# *Neisseria meningitidis* sibling small regulatory RNAs connect metabolism with colonization by controlling propionate use

**DOI:** 10.1101/2022.11.13.516309

**Authors:** Sandra Man-Bovenkerk, Kim Schipper, Nina M. van Sorge, Dave Speijer, Arie van der Ende, Yvonne Pannekoek

## Abstract

*Neisseria meningitidis* (the meningococcus) colonizes the human nasopharynx, primarily as a commensal, but sporadically causing septicemia and meningitis. During colonization and invasion, it encounters different niches with specific nutrient compositions. Small non-coding RNAs (sRNAs) are used to fine-tune expression of genes allowing adaptation to their physiological differences. We have previously characterized sRNAs (Neisseria metabolic switch regulators [NmsRs]) controlling switches between cataplerotic and anaplerotic metabolism. Here we extend the NmsRs regulon by studying methylcitrate lyase (PrpF) and propionate kinase (AckA-1) involved in the methylcitrate cycle, and serine hydroxymethyltransferase (GlyA) and 3-hydroxyacid dehydrogenase (MmsB) involved in protein degradation. These proteins were previously shown to be dysregulated in a Δ*nmsRs* strain. Levels of transcription of target genes and NmsRs were assessed by RT-qPCR. We also used a novel gene-reporter system, in which the 5’UTR of the target gene is fused to mcherry to study NmsRs-target gene interaction in the meningococcus.

Under nutrient-rich conditions, NmsRs downregulate expression of PrpF and AckA-1, by direct interaction with the 5’ UTR of their mRNA. Overexpression of NmsRs impaired growth under nutrient-limiting growth conditions with pyruvate and propionic acid as the only carbon sources. Our data strongly suggest that NmsRs downregulate propionate metabolism by lowering methylcitrate enzyme activity under nutrient-rich conditions. Under nutrient-poor conditions, NmsRs are downregulated, increasing propionate metabolism, resulting in higher tricarboxylic acid (TCA) activities. This allows metabolism to support nasopharynx colonization with breakdown products of amino acids functioning as anaplerotic substrates, as highlighted by NmsRs regulation of GlyA and MmsB.

**SIGNIFICANCE:** *Neisseria meningitidis* colonizes the human adult nasopharynx, forming a reservoir for the sporadic occurrence of epidemic invasive meningococcal disease like septicemia and meningitis. Propionic acid generated by other bacteria that co-inhabit the human adolescent/adult nasopharynx can be utilized by meningococci for replication in this environment. Here we showed that the sibling small RNAs designated NmsRs (Neisseria metabolic switch regulators) riboregulate propionic acid utilization by meningococci, and thus, colonization. Under conditions mimicking the nasopharyngeal environment, NmsRs upregulated expression of enzymes of the methylcitrate cycle. This leads to the conversion of propionic acid to pyruvate and succinate, resulting in higher tricarboxylic acid cycle activity, allowing colonization of the nasopharynx. NmsRs link metabolic state with colonization, which is a crucial step on the trajectory to invasive meningococcal disease.

## INTRODUCTION

*Neisseria meningitidis* (the meningococcus) normally resides innocuously in the nasopharynx. Only in rare cases can it become pathogenic, by passing the nasopharyngeal epithelium, entering and replicating in the bloodstream to cause septicemia, and subsequently crossing the blood-brain barrier to multiply in the cerebrospinal fluid (CSF), leading to meningitis. The sudden onset and progression of disease, often within only 24 h, contributes to a high incidence of mortality and morbidity (1). En route to bloodstream invasion, the meningococcus encounters different environmental conditions, comprising competing polymicrobial complexes, nutrients, temperature and/or relative gas concentrations, necessitating metabolic and protein repertoire adjustments. Its potential to adapt to changing conditions allow meningococci to survive and multiply during systemic infections (2).

The machinery of the meningococcus reorganizing cellular transcription (and thus gene expression) upon environmental challenges such as starvation and hypoxia, the so-called stringent response, is composed of regulatory transcription factors, two-component regulatory systems and small RNAs (sRNAs). Numerous sRNAs with sizes varying from 50 to 300 nucleotides were recently identified among a wide variety of pathogens. Many of these sRNAs function by imperfect anti-sense base pairing with multiple mRNA targets at or near the Shine-Dalgarno (SD) sequence of their targets, preventing ribosome-entry (3–5). This results in translational inhibition and is occasionally followed by degradation of the mRNA (6).

In general, only a relatively small number of sRNA targets have been functionally characterized, leaving most of these putative regulators with unknown functions. Interestingly, characterized sRNAs were involved in adaptation to varying environmental conditions such as encountered during the course of infection. They regulated the expression of colonization factors, toxins, and components involved in quorum sensing and evading host immune defenses: all virulence-related physiological and metabolic processes (7).

Previously, we identified and functionally characterized a novel class of sibling sRNAs (Neisseria metabolic switch regulators [NmsRs]) of meningococci. The sRNAs NmsR_A_ and NmsR_B_ share 70% sequence identity, are tandemly arranged and highly conserved in the genomes of all *Neisseria* species (8). NmsRs are expressed during high nutrient availability (e.g., within the bloodstream) but not in nutrient-depleted growth conditions such as present within the nasopharynx. Through antisense mechanisms, we showed that NmsRs downregulated the biosynthesis of four enzymes (SdhC, GltA, SucC and FumC), all involved in the TCA cycle, and two enzymes, methylcitrate lyase (PrpB) and methylcitrate synthase (PrpC) involved in methylcitrate cycle/propionate metabolism. Expression of NmsRs was shown to be controlled by the RelA-regulated stringent response, stressing their importance in surviving nutrient limitations and suggesting an important role in the switch from cataplerotic to anaplerotic metabolism (8).

The natural habitat of the meningococcus is the adolescent/adult nasopharynx, which is poor in nutrients but rich in anaerobe bacteria, many of which produce propionic acid (9, 10). To support its growth under nutrient-poor conditions, *N. meningitidis* can convert propionic acid into succinate and pyruvate, which are then used as a supplementary carbon source, via its propionate metabolism pathway (11). (**Fig. 1**, schematizing the TCA and the methylcitrate cycle of *N. meningitidis*). Use of propionic acid requires the *prp* gene cluster, which comprises the genes NMB0430-NMB435 in the serogroup B strain MC58 and is localized on a genomic island (11, 12). As mentioned, two enzymes of this cluster, PrpB/NMB0430 and PrpC/NMB0431, are under post-transcriptional control of NmsRs (8). Previously, we showed that NMB0434, coding for methylcitrate lyase (PrpF), NMB0435, encoding propionate kinase (AckA-1), NMB1055, encoding serine hydroxymethyltransferase (GlyA), NMB1584, encoding 3-hydroxyacid dehydrogenase (MmsB) (the latter two both involved in protein degradation), and a transamidase subunit involved in correct tRNA charging (GatC) are upregulated in a Δ*nmsRs* strain (8). We did not demonstrate whether this upregulation was a direct or indirect consequence of *nmsRs* deletion. To obtain further insight in the regulatory network of NmsRs and their significance for controlling general metabolic switches, we investigated whether PrpF, AckA-1, GlyA, MmsB and GatC are also part of the NmsR regulon. Moreover, we studied the biological significance of NmsRs-mediated regulation of expression of enzymes by involved in the methylcitrate cycle and, hence, propionate metabolism. Thus, we assessed the direct interaction between NmsRs and their potential mRNA targets and the regulatory effect of NmsRs on the growth kinetics of meningococci under growth conditions mimicking different *in vivo* conditions.

**Figure 1.**
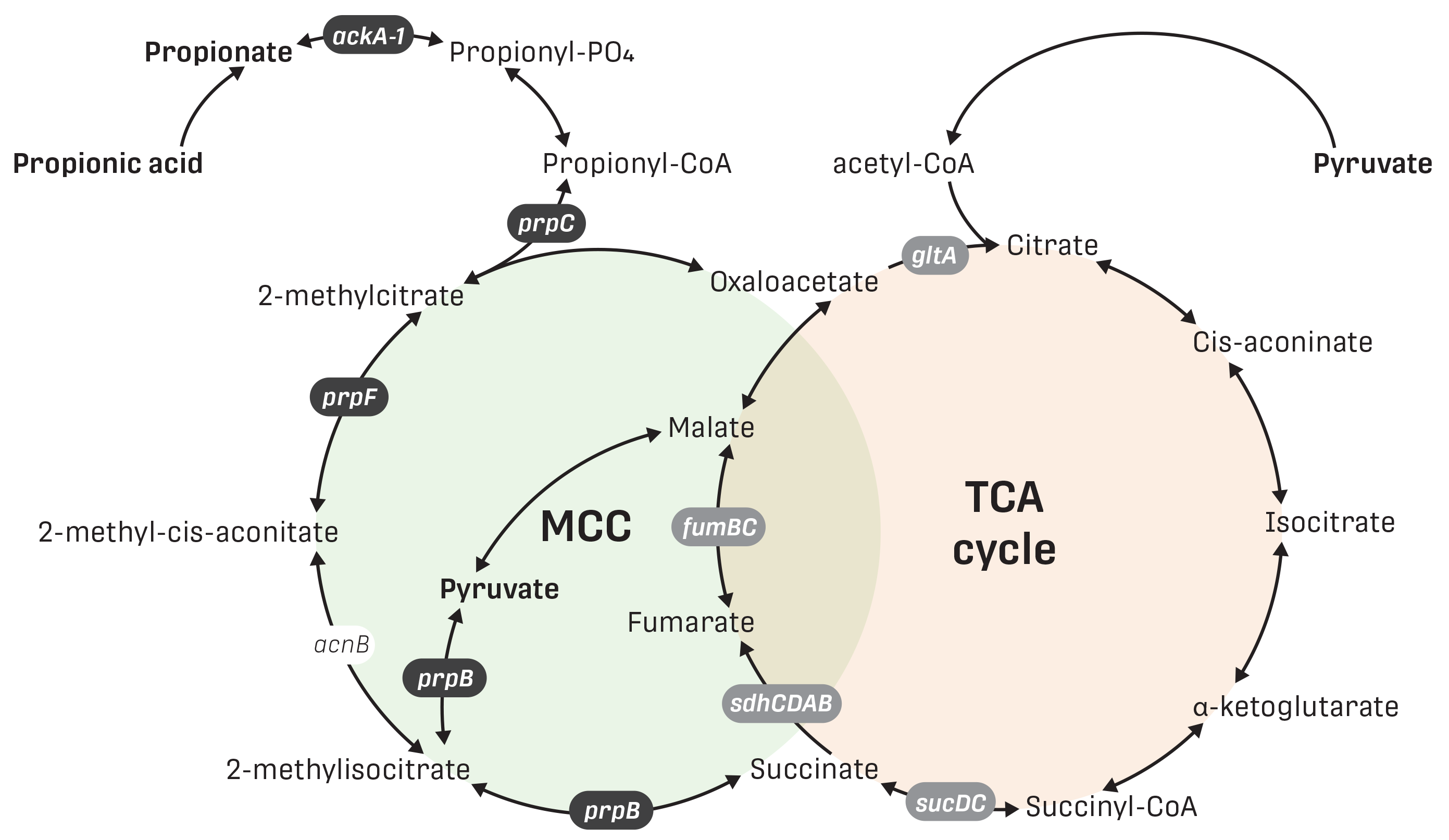
Schematic representation of the interactions between the MCC and TCA cycles. Overlapping reactions of methylcitrate (MCC) and the tricarboxylic acid (TCA) cycles in *N. meningitidis*. The substrates Propionic acid, Propionate and Pyruvate are indicated in bold. Previously demonstrated mRNA targets of NmsRs are shown in gray boxes.

## RESULTS

### NmsRs alter transcript levels of enzymes involved in the methylcitrate cycle and (branched chain) amino acid degradation

Proteomic analysis revealed higher enzyme levels of PrpF, AckA-1, GlyA, GatC and MmsB in meningococci upon *nmsRs* deletion (8). To investigate whether NmsRs directly interacted with the relevant transcripts, we first assessed their levels in meningococci grown under nutrient rich conditions. To this end, we created an *nmsRs* knock-out strain of H44/76 by replacing *nmsR_A_* and *nmsR_B_* with a Kanamycine (Kan) resistance cassette. Next, we complemented the *nmsRs* knock-out strain by introducing a plasmid harboring the both NmsRs-encoding genes. To correct for potential effects of plasmid-replication on strain characteristics, the empty plasmid was introduced into the wild type and the *nmsRs* knock-out strain, generating an isogenic panel of strains, consisting of wild type+pEN11 (wt), the *nmsRs* knock-out strain with pEN11 (Δ*nmsRs*) and Δ*nmsRs* overexpressing NmsRs (Δ*nmsRs*+*nmsRs*). Transcript levels of *prpF*, *ackA*-1, *glyA, mmsB*, and *gatC* were quantified by RT-qPCR. *PrpC* (encoding citrate synthase) and *porA* (encoding major outer membrane protein A), were used as NmsRs-down-regulated control (8) and NmsRs-non-regulated control (13), respectively. Compared to wt, transcript levels of *prpF, ackA*-1, *glyA, mmsB* and *prpC* were significantly higher (47- to 2-fold [*P*<0.05]) in Δ*nmsRs* meningococci, while being significantly lower (*P*<0.05) in the isogenic strain overexpressing NmsRs (**Fig. 2**). Transcript levels of *gatC and porA* were not significantly different between any of the strains (**Fig. 2**).

**Figure 2.**
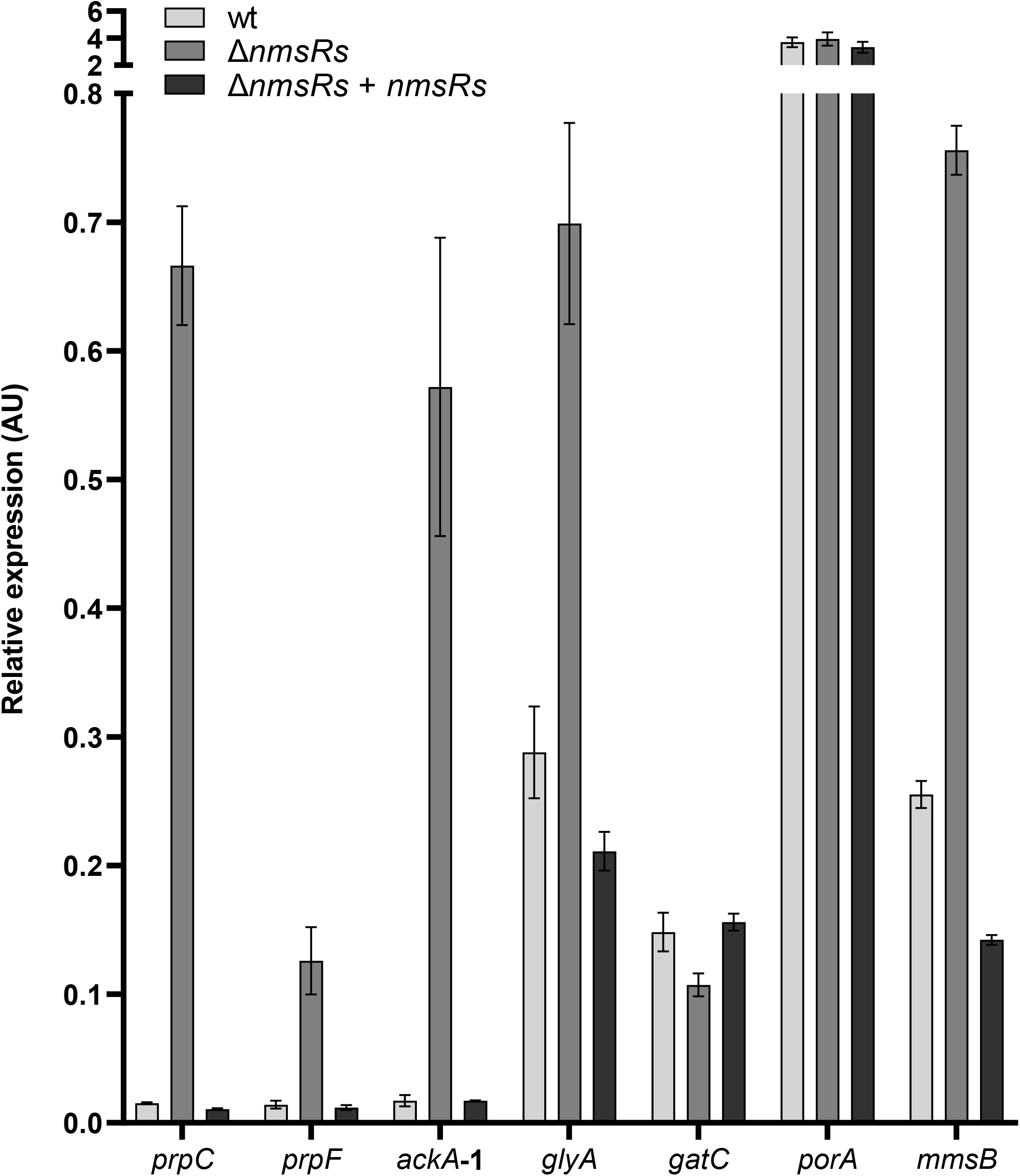
Transcript levels of *prpF*, *ackA*-1, *glyA* and *mmsB* are genetically linked to NmsRs. Transcript levels assessed by RT-qPCR in wt, Δ*nmsRs*, and a NmsRs overexpressing strain (Δ*nmsRs*+*nmsRs*) after growth in nutrient rich medium (error bars, standard errors of the means; technical replicates, n = 3, over biological, n = 4). *prpC* and *porA* function as positive and negative control, respectively.

### NmsRs translationally downregulate enzymes involved in propionate metabolism and (branched chain) amino acid degradation

*Trans*-acting sRNAs transcribed from intergenic regions usually have some complementarity (α-SDs regions; UC-rich) often in or near the 5’ UTR region of their target mRNA(s) that might allow partial sRNA-mRNA duplex formation. Duplex formation can either repress translation by obstructing the AG-rich SD region, or activate translation by preventing inhibitory secondary structures (masking the SD) of the mRNA itself (14–16). The NmsRs are predicted to fold into similar secondary structures consisting of three stem-loops (SLs) (**Fig. 3A**). The single stranded region between SL1 and SL2 exposes a UC-rich sequence, as does the single-stranded loop of SL2. These regions are (partly) complementary to the SD regions of previously proven targets of NmsRs (*prpB* and *prpC*) (**Fig. 3A**) (8).

**Figure 3.**
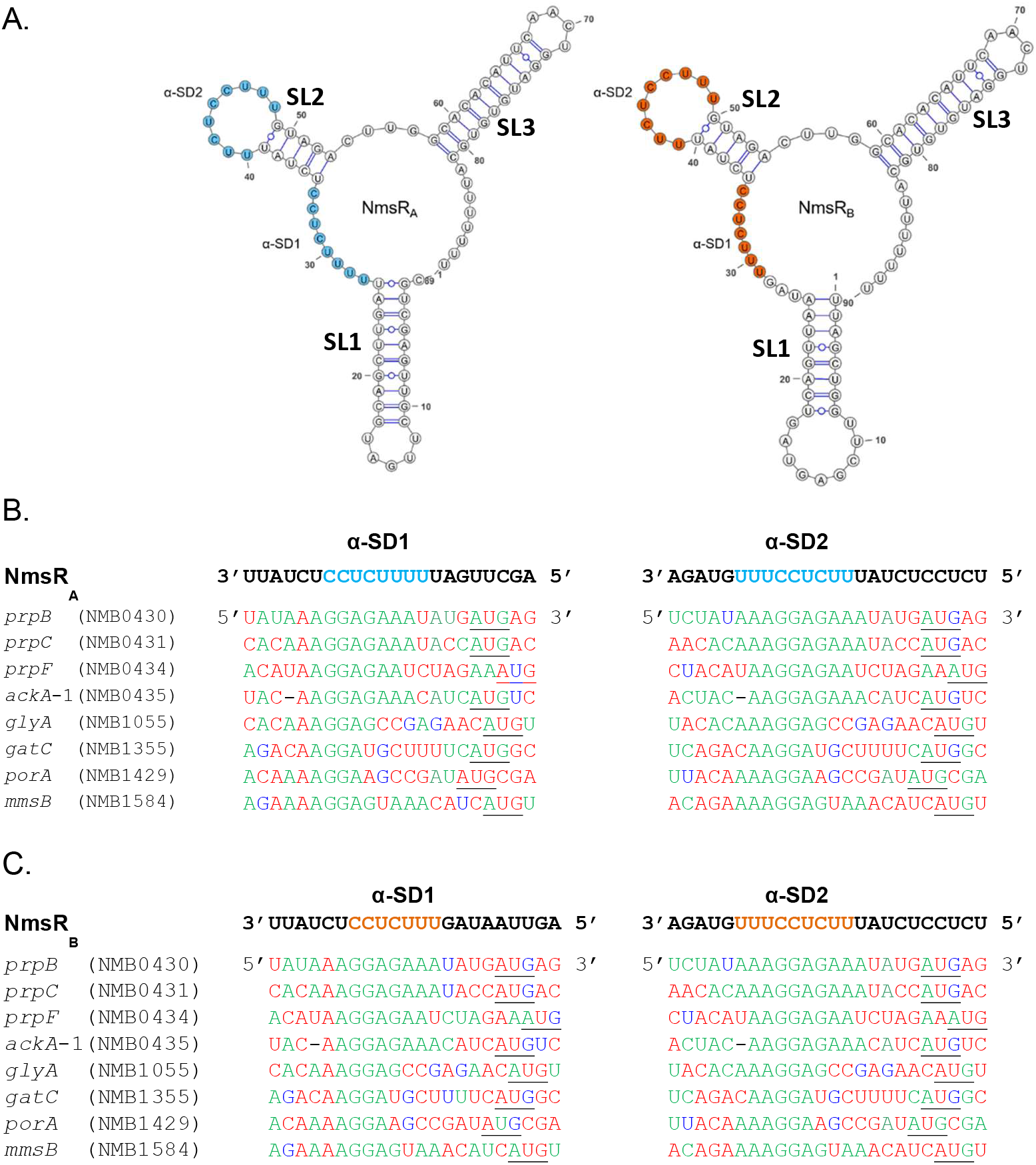
Schematic representation of the predicted secondary structures of NmsRs and interaction with the 5’ UTR regions of target mRNAs. (A) Predicted secondary structures of siblings NmsR_A_ and NmsR_B_ (minimum free energy of −30.9 and 22.0 kcal/mol, respectively). Stem loops are indicated as “SL”. Secondary structures were predicted using RNAfold (45) and visualized with VARNA (46) (Figure adapted from (10)). Putative α-SD sequences are circled blue in NmsR_A_ and orange in NmsR_B_. Nucleotides of 5′ UTRs of mRNAs predicted to be targeted by NmsR_A_ (B) and NmsR_B_ (C) are aligned with each other. Nucleotides complementary to NmsRs sequences are in green, nucleotides involved in possible G⋅U pairs in dark blue, mismatches in red, and AUG initiations codons are underlined.

We performed sequence analysis to investigate potential base-pair regions between the NmsRs and the 5’ UTR of the mRNA of *prpF, ackA*-1, *glyA, mmsB* and *prpC* (all with low transcription levels in the wt and Δ*nmsRs*+*nmsRs* strains). This revealed 5 to 7 bp complementarity with the 8 bp αSD1 of NmsR_A_ (**Fig. 3B**) and 4 to 7 bp complementarity with the 7 bp of αSD1 of NmsR_B_ (**Fig. 3C**). In contrast, the 5’ UTRs of *gatC* and *porA* (with similar transcription levels in wt, Δ*nmsR* and Δ*nmsRs*+*nmsRs*) showed complementarity stretches of only 3 bp with αSD1 of NmsR_A_ and NmsR_B_ respectively (**Fig 3. B and C**). Overall, the complementarity with the 9 bp of αSD2 of NmsR_A_ and NmsR_B_ varies between 7 and 9 bp for *prpF, ackA*-1, *glyA, mmsB* and *prpC* and 5 and 6 bp for *gatC* and *porA*, respectively.

To obtain further experimental evidence for interactions between the NmsRs and the candidate mRNA targets and insight in the mechanism of regulation, we developed a novel reporter system meningococci to assess post-transcriptional regulation of targets mRNAs. Therefore we used plasmid pMGC5’ (17) containing the constitutive *pilE* promoter and generated translational fusions between the 5’ UTR and the first 15 to 18 codons of putatively regulated genes in frame to *mcherry* (*gene*∷*mcherry*), downstream of this promoter. Constructs were integrated in the chromosome of Δ*nmsRs* by homologous recombination, thereby generating Δ*nmsRs*+*gene*∷*mcherry* strains (**Fig. 4A)**. Next, Δ*nmsRs*+*gene*∷*mcherry* strains were transformed with either pEN11 or pEN11_NmsRs to generate isogenic strains carrying the translational fusions, with (Δ*nmsRs*+*gene*∷*mcherry*+*nmsRs*) or without (Δ*nmsRs*+*gene*∷*mcherry*) expression of NmsRs. Reduced fluorescence in the meningococcus in the presence of NmsRs (Δ*nmsRs*+*gene*∷*mcherry*+*nmsRs*) compared to the isogenic strain without NmsRs, indicates translational downregulation by direct interaction between NmsRs and 5’ UTRs of their target RNAs. For validation, PrpC was used as NmsRs-downregulated control (8) and PorA as nonregulated control (13).

**Figure 4.**
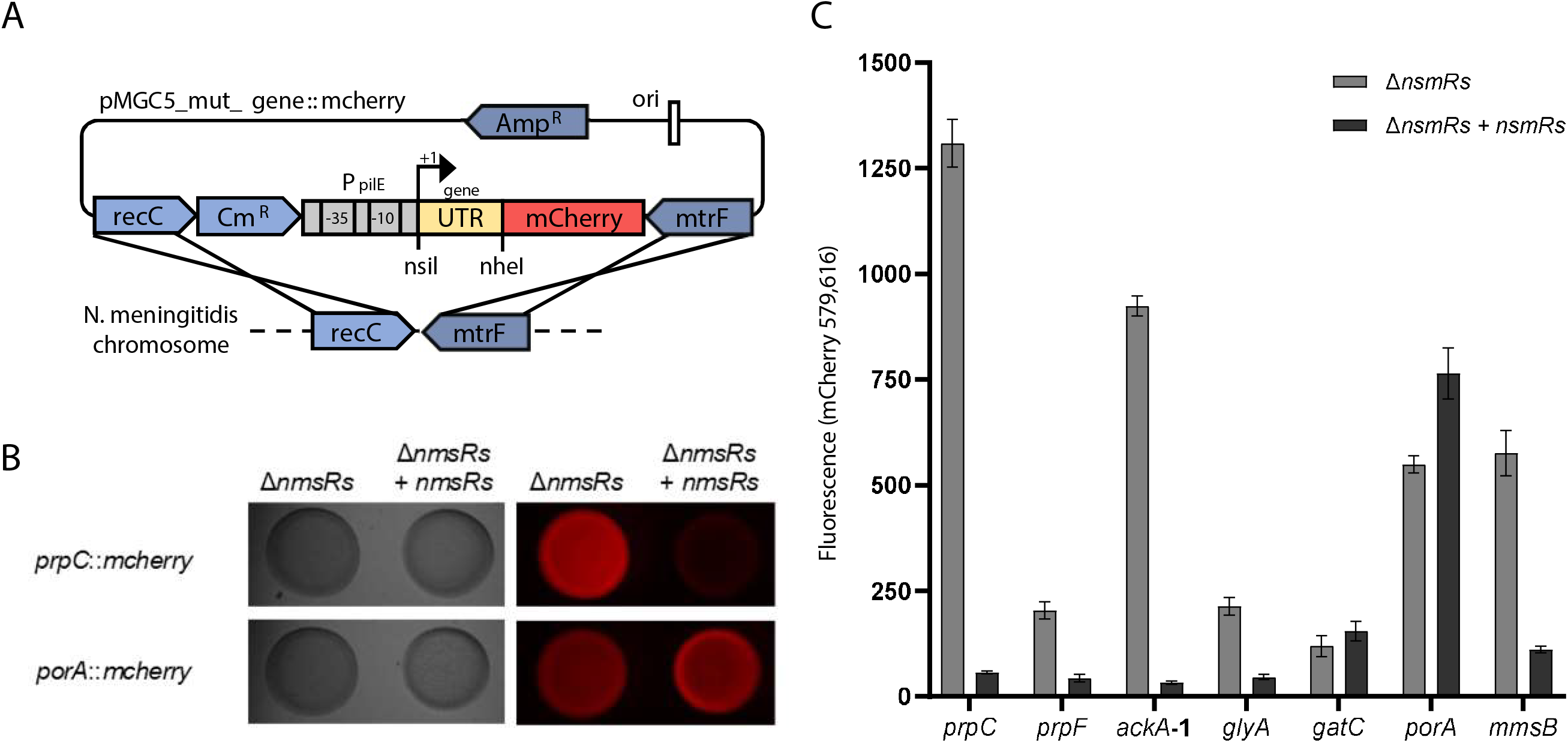
Translational down regulation of target mRNA expression by NmsRs. (A) Schematic illustration of the integrational vector pMGC5_mut_*gene*∷*mcherry* used for construction of translational *mcherry* fusions to be inserted into the *recC* and *mtrF* locus of *N. meningitidis* H44/76. (47). The 5 ’UTRs and the first 15 to18 amino acids of the coding regions (yellow box) of candidate targets were fused to the fluorescence *mcherry* reporter gene (red box), transcribed from the constitutive *pilE* promoter as indicated by the arrow. Elements are not drawn to scale. (B) Images of GC agar plates of meningococci carrying chromosomes with the translational fusion *prpC*∷*mcherry* or *porA*∷*mcherry*, with (Δ*nmsRs*) or without (Δ*nmsRs*+*nmsRs*) NmsRs. Images obtained with visible light (left) or in fluorescence mode (right). Reduced fluorescence of the *prpC*∷*mcherry* fusion strain upon NmsRs co-expression indicates regulation at the post-transcriptional level. (C) Regulation of translational fusions of *prpC*∷*mcherry*, *prpF*∷*mcherry*, *ackA*-1∷*mcherry*, *glyA*∷*mcherry* and *mmsB*∷*mcherry*. Quantification of specific fluorescence in cells carrying the translational *gene*∷*mcherry* fusions in the chromosome, without (Δ*nmsRs*) or with NmsR (Δ*nmsRs+nmsRs)* present, upon growth in nutrient rich medium (error bars, standard errors of the means; technical replicates, n = 3, over biological, n = 3).

As predicted, fluorescence levels of Δ*nmsRs* meningococci carrying the translational *prpC∷mcherry* fusion and overexpressing *nmsRs* were 25-fold lower (*P* < 0.005) than those of Δ*nmsRs* carrying the translational *prpC*∷*mcherry* fusion without NmsRs. (**Fig. 4B and C)**. Fluorescence levels of cells containing *porA*∷*mcherry* were indeed not significantly different between Δ*nmsRs* and Δ*nmsRs*+*nmsRs* cells (**Fig. 4B and C**).

Direct translational downregulation by NmsRs was demonstrated for four out of five tested putative targets in the *mcherry*-reporter system (*prpF*, *ackA*-1, *glyA* and *mmsB* (*P* < 0.05) (**Fig. 4C)**. Fluorescence levels of *gatC* were comparable with or without NmsRs, ruling out direct NmsRs-mediated regulation (**Fig. 4C**).

### NmsRs impair extended growth in medium with propionic acid as a carbon source

Since NmsRs-regulated transcripts are involved in the methylcitrate cycle, we further explored the biological significance of NmsRs-regulation of propionate metabolism. Growth characteristics of meningococci were investigated in nutrient-limited medium (18) with pyruvate and propionic acid as sole carbon sources.

In medium supplemented with pyruvate only, both the wt and Δ*nmsRs* variant replicated equally well and reached a maximum of an OD_600_ of ~ 0.5 in 6 h to 7 h (**Fig. 5A and B**, open symbols). Growth was extended for another 3 h when propionic acid was also added to the medium, reaching an OD_600_ of 0.6 to 0.7. However, growth of the NmsRs overexpressing strain was slower in both media and only reached an OD_600_ of ~ 0.4 after 6 h of growth (**Fig. 5C**). Thus, overexpression of NmsRs results in the inhibition of extended growth in medium with propionic acid. This suggest that NmsRs interfered with the utilization of propionic acid, which starts around 4-5 h (11), presumably by lowering propionate-processing enzyme concentrations.

**Figure 5.**
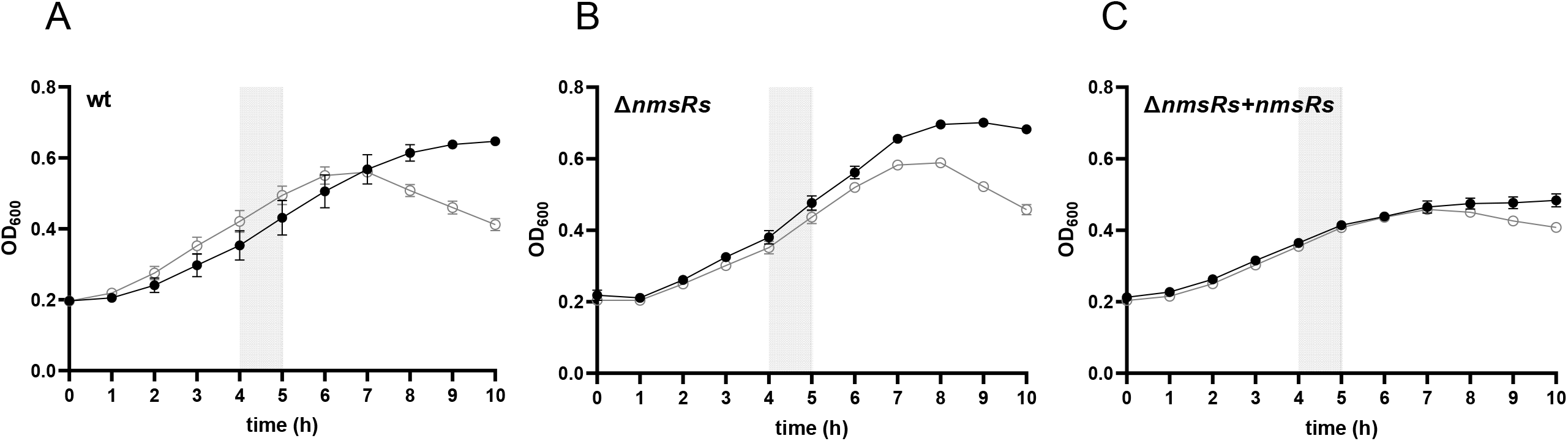
Overexpression of NmsRs leads to a growth defect in minimal medium with pyruvate and propionic acid as sole carbon sources. Growth characteristics of different meningococcal strains. (A): wildtype (wt); (B): meningococci with *nmsRs* deleted (Δ*nmsRs*); (C): meningococci overexpressing NmsRs (Δ*nmsRs*+*nmsRs*). Growth in minimal medium with pyruvate only (10 mM) (open symbols) and pyruvate (10 mM) and propionic acid (5 mM) (filled symbols). Points represent means from technical replicates n = 3, over biological experiments, n = 3 (error bars, standard errors of the means). The grey area indicates the approximately time frame of start of propionic acid utilization (11).

### NmsRs downregulate enzymes of the *prp* gene cluster

To confirm that NmsRs lowered propionate-processing enzymes, we analyzed expression of *prpB*, *prpC*, *prpF* and *ackA*-1 during growth in minimal medium with propionic acid after pyruvate is depleted.

Transcript levels of genes of the *prp* gene cluster, as well as NmsRs transcript levels themselves, were compared before (3 h) and after (9 h) pyruvate depletion in the absence and presence of propionic acid, using quantitative RT-qPCR analyses. After 3 h of growth, transcript levels of *prpB*, *prpC*, *prpF* and *ackA*-1 of wt cells grown in medium with pyruvate only were not significantly different from those in bacteria grown in medium with pyruvate and propionic acid (**Fig. 6A**). In contrast, after 9 h of growth, transcript levels of *prpB*, *prpC*, *prpF* and *ackA*-1 were significantly (3.6- to 8.2-fold) higher in meningococci grown in medium with pyruvate and propionic acid compared to medium with pyruvate only (*P* < 0.01) (**Fig. 6A**).

**Figure 6.**
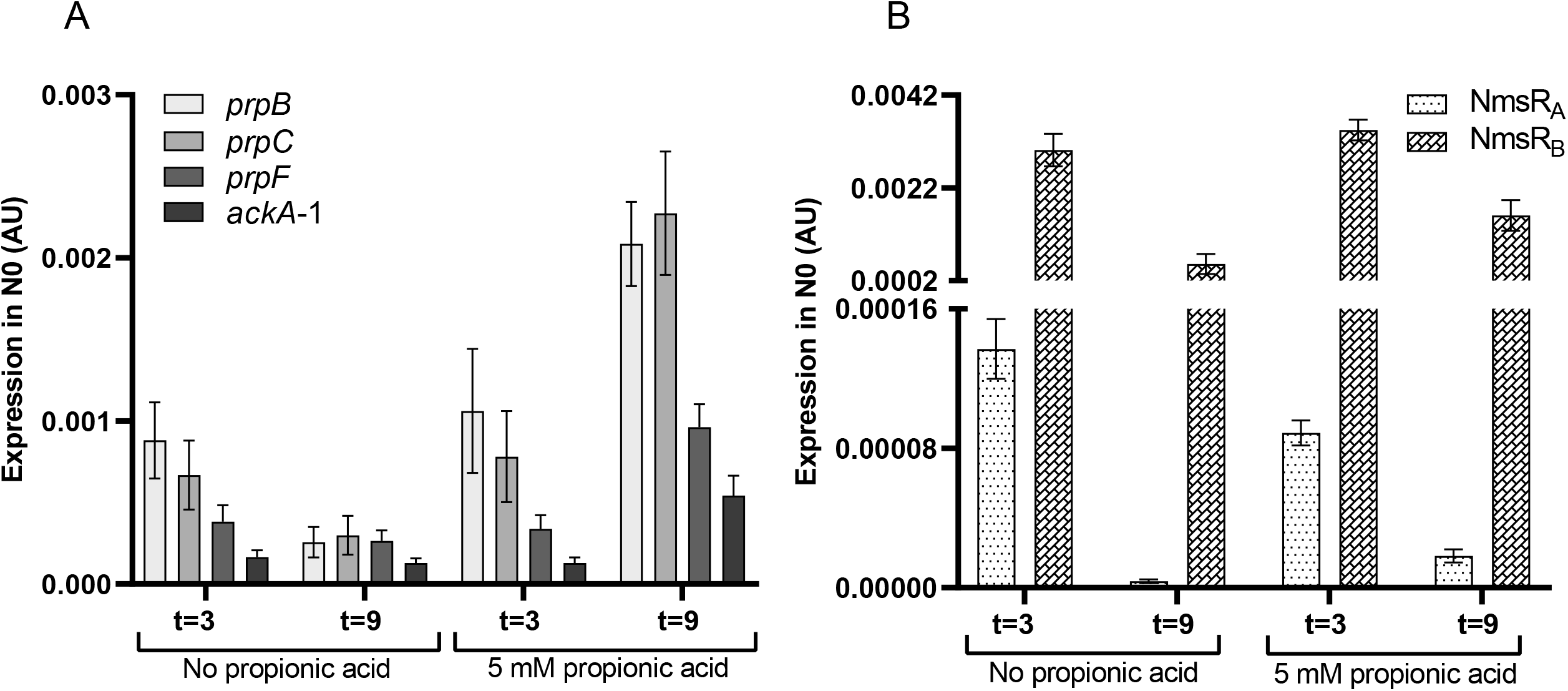
Transcript levels of wt meningococci during growth in minimal medium with pyruvate and varying amounts of propionic acid as sole carbon sources. Transcript levels assessed by RT-qPCR in wt meningococci (error bars, standard errors of the means; technical replicates, n = 2, over biological experiments, n = 3). Meningococci were cultured in minimal medium with pyruvate only, or with pyruvate and propionic acid. Transcript levels of (A) *prpB*, *prpC*, *prpF* and *ackA*-1 and (B) NmsRs were assessed after 3 and 9 h growth.

After 3 h and 9 h of growth, transcript levels of NmsRs in wt cells were not significantly different in medium with pyruvate only or in medium with pyruvate and propionic acid (**Fig. 6B**). In contrast, after 9 h of growth, transcript levels of NmsRs were significantly lower (2.1- to 4.9-fold in the presence of propionic acid) compared to levels after 3 h growth under both conditions (*P* < 0.01) (**Fig. 6B)**.

Of note, transcript levels of the four target genes in Δ*nmsRs* cells after 3 h of growth were significantly (3.7 to 12.6-fold [*P* < 0.05]) higher compared to wt (**Fig. S1**). Except for *ackA*-1, transcript levels of the target genes in Δ*nmsRs* cells were significantly (3.2 to 1.8-fold [*P* < 0.008]) higher after 9 h of growth in medium with propionic acid compared to wt cells. (**Fig. S1**). In the strain overexpressing NmsRs, NmsR_B_ levels were significantly higher (16.6 to 17.1-fold [*P* < 0.0001]) compared to those of wt **(Fig. S2)**, while no transcripts were detected in Δ*nmsRs* (not shown).

Taken together, our results strongly indicate that expression of NmsRs results in downregulation of expression of the genes of the *prp* cluster assessed, most likely by post-transcriptional regulation resulting in translational repression. This is accompanied by slower growth with propionic acid as sole carbon source after pyruvate depletion.

## DISCUSSION

Here we provide further insight in the regulatory network of NmsRs and its control of metabolic switches in *N. meningitidis*. Our data show regulation of the methylcitrate cycle and propionate metabolism by direct NmsRs actions. This activity is crucial when meningococci encounter different host environments, varying in nutrients, like the adolescent/adult nasopharynx containing propionic acid as a supplementary carbon source (9).

Using a well-established *gfp*-reporter system in *E. coli* (19), we previously demonstrated translational repression of *prpC* and *prpB* mediated by NmsRs (8). Now we show, using a newly developed *mcherry*-reporter system in the genetic background of meningococci, translational repression of other *prp* gene cluster genes (*prpF* and *ackA*-1) by NmsRs. Besides these gene cluster members, we also demonstrated NmsRs-mediated translational repression of serine hydroxymethyltransferase (GlyA) and 3-hydroxyacid dehydrogenase (MmsB). These proteins function in branched chain amino acid (BCAA) degradation (Ile, Leu, and Val). Importantly, in contrast to *prpC*∷*mcherry* (positive control), the *porA*∷*mcherry* (negative control) was not affected by NmsRs, validating the system and ruling out non-specific effects of mCherry and/or NmsRs expression.

These novel NmsRs targets, as well as GatC, were known to be dysregulated in a Δ*nmsR* strain (8). Sequence analysis of the degree of complementarity between the UC-rich α-SD regions of NmsRs and the AG-rich regions of the 5’ UTRs of the novel mRNA targets revealed relatively long stretches of complementarity ranging from 5-6/8 or 7/8 nucleotides and 4/7 to 7/7 nucleotides with the α-SD1s of NmsR_A_ and NmsR_B_, respectively. The 5’ UTRs of *gatC* and *porA* showed a stretch of complementarity of only 3 nucleotides long. Similarly, the complementarity between the 5’ UTR of *porA* and *gatC* with the α-SD2 of NmsR_A_ and NmsR_B_ was smaller than that between the other targets and the both α-SD2s (5/9 or 6/9 vs 7/9 to 9/9). The expression of *porA* and *gatC* was shown to be independent of NmsRs in the meningococcal reporter system, consistent with transcript analyses, showing their transcript levels remained comparable to wt upon deletion or overexpression of NmsRs. This might point to minimal requirement for duplex formation between NmsRs and the 5’ UTRs of target genes, being larger than 3 bp and 5 to 6 bp for α-SD1 and α-SD2, respectively. Upregulation of GatC in Δ*nmsR*, found in proteomic analysis of Δ*nmsRs* cells, might indicate an indirect effect upon expression of *gatC* and suggests that regulation of this gene is more complex, e.g. controlled by other regulators connected to the NmsRs regulon.

Translational repression, as monitored with our reporter system, is often coupled to destabilization of the respective mRNAs due to the connection between ribosome loading and mRNA stability (14–16). sRNA binding may also directly affect mRNA levels by recruiting RNA degrading enzymes (RNaseE of RNAaseIII) leading to rapid degradation of the mRNA (14–16). Upon overexpression of NmsRs in the meningococcus, significant lower transcript levels of all assessed novel targets (excepting *gatC*) were observed, indicating transcript levels of the target mRNAs are rapidly degraded upon duplex formation with NmsRs. However, the precise mechanisms and nucleases involved in such processes in *N. meningitidis* await further studies. Also the role of meningococcal RNA chaperones, involved in facilitating the interaction between sRNAs and their target mRNAs has to be elucidated (20). The Hfq chaperone and FinO-domain proteins like ProQ represent important examples, regulating numerous complex processes, including bacterial growth, the stringent response and virulence (21–23). All NmsRs targets of our study were also overexpressed in a H44/76 Hfq deletion strain (24) but apparently not in a meningococcus without ProQ (strain 8013; serogroup C) (25). The latter observation could be indicative of distinct targetomes of Hfq and ProQ in meningococci as previously suggested (25), but might be more complicated as well (26, 27).

Deletion of *relA* results in elevated NmsRs levels suggesting that expression of NmsRs is (in)directly controlled by (p)ppGpp levels (8). Enzymes degrading BCAAs (Ile, Leu, and Val), such as GlyA and MmsB, are considered biological markers of the stringent response (26). In addition, BCAAs are also effectors of global transcriptional regulators such as the leucine-responsive regulatory protein (Lrp) and the RelA (27, 28). These enzymes therefore could reflect a physiological state of the meningococcus restructuring protein/amino acid metabolism. The observation that expression of these enzymes is also regulated by NmsRs confirms the connection between the stringent response and the NmsRs regulated network.

Among the very few sRNAs in *N. meningitidis*, the blood-induced sRNA Bns1 is of particular interest in the context of the present study (29). The predicted target mRNA of Bns1 is NMB0429, which is located within the *prp* cluster (30). In addition, Bns1 is subjected to negative regulation by GntR/NMB1563 in the absence of carbon sources (30). Furthermore, it has been reported that in meningococci *prpB*, *prpC* and *acnA* were upregulated in meningococci in which *misR* - part of the two-component system *misR*/*mirS* (NMB0595/0594) - was deleted (31). Although in none of the aforementioned studies direct regulation of genes of the *prp* gene cluster by Bns1 or MisR was assessed, these candidate regulators could combine with NmsRs in complicated layers of *prp* gene cluster regulation in the meningococcus. This could explain that after 9 h transcript levels of the studied *prp* genes were significantly lower in cultures when propionic acid was not added (**Fig. 2A, Fig. S2A**), while NmsRs levels were not significantly different in both growth conditions after 9 h (**Fig. 2B**). Together, this might indicate that in meningococci, transcription of the *prp* gene cluster is not solely controlled by NmsRs but also by as yet unknown regulator(s), repressing in the absence and/or activating in the presence of propionic acid-derived stimuli.

Of note, gonococcal counterparts of NmsRs (NgncR_162 and NgncR_163) were also shown to interact with *prpB*, *prpC* and *ackA*-1, while controlling the transcriptional regulator GdhR as well (32). In gonococci, GdhR directly, negatively, regulates *lctP*, encoding an L-lactate permease required for successfully colonization of the murine genital tract (33). Recently, lactate uptake has been implicated in resistance to neutrophils (34). The relevance of NgncRs in propionic acid metabolism remains unclear, but just as NmsRs manipulate propionic acid metabolism beneficial for nasopharynx colonization, NgcRs might be manipulating lactate metabolism to colonize the female reproductive tract (by regulating GdhR). In meningococci, GdhR/NMB1711 mediates growth phase- and carbon source dependent positive regulation of *gdhA*, coding for L-glutamate dehydrogenase, essential for systemic infection in an infant rat model (35, 36). However, in gonococci *gdhA* was not found to be under control of GdhR, suggesting that the regulons do not share similar functions, likely due to differences in regulatory sequences in promoter regions (37). Our preliminary data showed significant (*P* < 0,05) translational repression of GdhR in the gene-reporter system in meningococci (not shown). However, transcript levels were not significantly different between wt, Δ*nmsRs* or a strain overexpressing NmsRs (not shown). Whether GdhR is also under control of NmsRs awaits further studies, but if *gdhR* mRNA is a true target of NmsRs, than its interaction with NmsRs is not directly coupled to its destabilization (as found for all other known mRNA targets of NmsRs). This could point to different mechanisms of translational repression.

During growth of wt meningococci in minimal medium with pyruvate we observed growth retardation after 6 h, but in the presence of propionic acid extended growth. Overexpression of NmsRs resulted in retarded growth at an earlier time point (4-5 h), irrespective of the presence of propionic acid. During growth in minimal medium supplemented with pyruvate and propionic acid, propionic utilization already starts after 4-5 h (11). Apparently under these growth conditions a nutrient limiting environment (stringent response) is encountered, as exemplified by the utilization of propionic acid. This explains why overexpression of NmsRs, and subsequent downregulation of some of the *prp* cluster genes might inhibit growth early (after 4-5 h) on. In *N. meningitidis*, the genes of the methylcitrate cycle gene cluster (*prpB*, *prpC, prpF* and *ackA*-1) are located on a large genomic island. This genomic island is absent from closely related non-pathogenic *Neisseria* species such as *Neisseria lactamica* (11). The methylcitrate cycle gene cluster enables meningococci to convert propionic acid to pyruvate, thereby supporting growth as well as limiting the toxicity of propionic acid. Since the adolescent/adult nasopharynx is rich in propionic acid generating bacteria, the ability to utilize propionic acid is suggested to confer a colonizing advantage on meningococci (11). Importantly, the ability of meningococci to effectively colonize the adolescent/adult nasopharynx plays a crucial role in the transmission and epidemiology of meningococcal disease (38). To conclude, we identified novel targets of NmsRs, coding for enzymes involved in the methylcitrate cycle/propionate metabolism as well as for enzymes involved in BCAA degradation. The NmsRs regulon encompasses mRNA targets coding for genes functioning in the adaptation of the meningococcus to a wide variety of growth conditions reflecting changing host environments. The regulation of genes involved in propionate metabolism adds another level of complexity to the network of NmsRs and connects metabolic status with colonization, impacting virulence.

## MATERIALS AND METHODS

### Bacterial strains and culture conditions

All bacterial strains and plasmids used in this study are listed in **Table S1**. Meningococci were grown for 16-18 h on GC plates (Difco) supplemented with 1% (vol/vol) Vitox (Oxoid) at 37°C in a humidified atmosphere of 5% CO_2_. Broth culturing was performed on a gyratory shaker (200 rpm) at 37°C in GC broth supplemented with 1% (v/v) Vitox, or in modified Jyssum medium (18) in which glucose was replaced with pyruvate (10 mM), or with both pyruvate (10 mM) and propionic acid (5 mM). When appropriate, GC was supplemented with erythromycin (Erm) (5 μg/ml), and/or chloramphenicol (Cm) (10 μg/ml), and/or kanamycin (Kan) (50 μg/ml). Growth was monitored by measuring the optical density of cultures at 600 nm (OD_600_) at regular intervals. *Escherichia coli* was grown in Lysogeny broth composed of 1% tryptone (w/v), 0.5% yeast extract (w/v) and 1% NaCl (w/v) (LB) or on LB agar (1% [w/v]) plates at 37°C. When appropriate, LB was supplemented with Cm (10 μg/ml). Plasmid pMGC5 was used for cloning *gene*∷*mcherry* constructs (17). Shuttle vector pEN11 (39) was used to express sRNAs in meningococci. Plasmid DNA from *E. coli* was isolated from overnight cultures grown in LB using the GeneJET plasmid miniprep Kit (Thermo Fisher). DNA was gel purified using a GeneJET gel extraction kit (Fermentas). Digestion and ligation were carried out using enzymes supplied by Thermo Scientific. PCR was performed using a Thermo PCR machine (Biometra Trio, Analytic Jena). *N. meningitidis* was transformed as described previously (40). Transformants were plated on selective plates containing appropriate antibiotics and checked for correct integration by PCR. All constructs were verified by Sanger sequencing.

### Construction of Δ*nmsRs* of *N. meningitidis* strain H44/76 and overexpression of *nmsRs*

All oligonucleotides used in this study are listed in **Table S2**. *N. meningitidis* H44/76 knockout mutant of *nmsR_A_ and nmsR_B_* (Δ*nmsRs*) was constructed using a Gblock gene fragment (IDT) consisting of a fragment of the gene upstream of *nmsR_A_* (*dsbB*, NMB1649), a Kan resistance cassette (41) and a fragment of the gene downstream of *nmsR_B_ (lrp*, NMB1650). This fragment was amplified and transformed to H44/76 WT to generate the knockout strain (Δ*nmsRs*). The construction of plasmid overexpressing *nmsRs* (pEN11_NmsRs) in meningococci has been previously described (8, 42). H44/76 overexpressing NmsRs (Δ*nmsRs*+*nmsRs*) was created by transformation of Δ*nmsRs* with pEN11_NmsRs. As control Δ*nmsRs* was also transformed with pEN11.

### Construction of a *mcherry*-based translational fusion reporter system in *N. meningitidis*

To allow validation of predicted NmsRs targets we developed a *mcherry*-based translational fusion reporter system using the backbone of plasmid pMGC5 carrying *mcherry* under control of the *pilE* promoter to allow chromosomal expression in the genetic background of the meningococcus (17). We introduced an NsiI restriction site downstream of the *pilE* promoter at the transcriptional start site (+1) and an NheI restriction site directly after the ATG start codon of *mcherry* in pMGC5 by site-directed mutagenesis (Stratagene), creating plasmid pMGC5_mut. For cloning of *mcherry* fusions in pMGC5_mut, chromosomal fragments of H44/76 were amplified by PCR with a sense oligonucleotide annealing to the +1 (43) of the putative target gene of NmsRs and adds an NsiI restriction site and an antisense oligonucleotide which annealing to the N-terminal coding region of the putative target gene, while adding an in frame NheI restriction site. Typically, the full-length 5’ UTR (from +1 of the most proximal promoter) and the first 15-18 codons of the N-terminal region of the gene were digested with NsiI and NheI and cloned in NsiI-NheI pre-digested pMGC5_mut, thereby creating a translational fusion in pMGC5_mut_*gene*∷*mcherry*. These constructs were transformed to *E. coli* TOP10. After overnight growth, colonies were checked for fluorescence in a Syngene Bio Image analyzer using a Lumenera camera with excitation at 579 nm and a 616-nm emission filter. Brightly fluorescent Cm-resistant clones were selected and further propagated in *E. coli* TOP10 cells.

Next, *gene*∷*mcherry* fragments were integrated in the *recC*/*mtrF* locus of chromosome of Δ*nmsRs* meningococci by natural transformation of pMGC5_mut_*gene*∷*mcherry*, creating translational fusions in the genetic background of Δ*nmsRs* (Δ*nmsRs+gene*∷*mcherry*) (**See Fig. 4A** for a schematic illustration of the integrational vector). All Δ*nmsRs+gene*∷*mcherry* variant strains were transformed with pEN11 as well as with pEN11_NmsRs.

### Fluorescence measurements of *gene*∷*mcherry* translational fusions and data processing

Cell suspensions of Δ*nmsRs* variant strains expression *gene*∷*mcherry* fusions were diluted to OD_600_ of 0.1 and droplets (3 μl) were spotted on GC plates containing appropriate antibiotics. After overnight growth, spots were photographed using a M205FA microscope (Leica) with excitation at 579 nm and a 616-nm emission filter. Images were analyzed using the Leica application Suite advanced Fluorescence software. To measure whole-cell fluorescence in broth cultures, *gene*∷*mcherry* fusions transformed with pEN11 or pEN11_NmsRs were cultured overnight on plates and resuspended into GC broth at an OD_600_ of 0.1 in 20 ml GC broth in 100 ml sterile bottles (in triplicates). Cultures were grown for 3 h (mid logarithmic phase; linear range of increasing fluorescence). Then, triplicate aliquots of each culture were diluted to an OD_600_ of 0.2 and transferred to a 96-well microtiter plate (Greiner). Fluorescence was measured using a Biotek microplate reader (Synergy H1) with excitation at 579 nm and a 616-nm emission filter. Auto-fluorescence of the GC medium was subtracted from the samples to obtain specific fluorescence.

### RNA isolation, quantitative RT-PCR (RT-qPCR) and data processing

Total RNA was extracted from meningococci using the Direct-zol RNA miniprep kit (Zymo Research) followed by a DNase treatment using the Turbo DNA-free kit (Invitrogen). Then, cDNA was synthesized from 2 μg of RNA and random oligonucleotide hexamers using ThermoScript RT-PCR System for First-Strand cDNA Synthesis (Invitrogen). Absence of genomic DNA was controlled by no-RT reactions. Quantitative PCR was performed in triplicate (or more), on cDNA samples using the LightCycler 480 SYBR Green I Master kit in the CFX384 RT-PCR Detection System (Biorad). The identities of the resulting amplicons were checked by melting-curve analysis using the CFX384 software. Reaction mixtures containing no template were included in each experiment to check for contamination. Transcripts of target and reference genes (*rmpM* [NMB0382] and *cbbA* [NMB1869]). were analyzed using LinRegPCR for calculating the N_0_ starting concentration per sample (44). Constitutive relative gene expression was determined as a ratio of the target gene to the geometric means of the reference genes. Data are expressed as relative expression in arbitrary fluorescence units. If normalization was not allowed due to transcriptional variation of reference genes data are expressed as N_0_ values as indicated. The regulatory effect of NmsRs was also calculated as fold regulation.

### Statistical analyses

An unpaired *t* test was used to determine significant difference (*P* ≤ 0.05) of specific fluorescence between the variant Δ*nmsRs+gene*∷*mcherry* strains transformed with pEN11 (unregulated *gene*∷*mcherry* specific fluorescence) and pEN11_NmsRs (regulated *gene*∷*mcherry* specific fluorescence). The regulatory effect of NmsRs on a *gene*∷*mcherry* fusion was also expressed as fold-regulation (mean of three biological replicates). This was calculated by dividing the unregulated (Δ*nmsRs+gene*∷*mcherry* + pEN11) specific fluorescence by the regulated (Δ*nmsRs+gene*∷*mcherry* + pEN11_NmsRs) specific fluorescence. If a change of ≥ 1.5-fold up or down regulation was found, a *t* test was performed to determine significance (*P* ≤ 0.05).

Technical replicates (n=2 or n=3) of biological experiments (n=3 or n=4) where used for statistical analysis of transcript levels. One-way ANOVA Tukey’s multiple comparison tests were performed for assessing the statistical significance of the difference between means of the (normalized) N_0_ values.

## ACKNOWLEDGMENTS

We wish to thank B. van der Steen for his technical assistance and dr. W.H. Man for his assistance in shaping figures.

---

**Figure S1. Transcript levels of *prpB*, *prpC*, *prpF* and *ackA*-1 are under control of NmsRs**. Relative transcript expression levels of mRNA targets of NmsRs in different variant meningococcal strains. (A) wt; (B) meningococci in which the *nmsRs* have been deleted (Δ*nmsRs*); (C) meningococci overexpressing NmsRs (Δ*nmsRs*+*nmsRs*). Growth in minimal medium with pyruvate and propionic acid as sole carbon sources. Transcript levels assessed by RT-qPCR (error bars, standard errors of the means; technical replicates, n = 2, over biological experiments, n = 3). Meningococci were cultured in minimal medium with pyruvate only or with pyruvate and propionic acid and transcript levels of *prpB*, *prpC*, *prpF* and *ackA*-1 were assessed after 3 and 9 h growth.

**Figure S2. Transcript levels of NmsR_A_ and NmsR_B_ during growth in minimal medium with pyruvate and with or without propionic acid as sole carbon sources**.

Transcript levels of NmsR_A_ (A) and NmsR_B_ (B) in wt and meningococci overexpressing NmsRs (Δ*nmsRs*+*nmsRs*) during growth in minimal medium with pyruvate and propionic acid as sole carbon sources. Transcript levels assessed by RT-qPCR (error bars, standard errors of the means; technical replicates, n = 2, over biological experiments, n = 3). Meningococci were cultured in minimal medium with pyruvate only or with pyruvate and propionic acid and transcript levels of NmsRs were assessed after 3 and 9 h growth. Transcript levels in Δ*nmsRs* were not detectable (not shown).

**Table S1. Strains and plasmids used in this study**.

**Tabel S2. Oligonucleotides used in this study**.

Sequences are given in the 5’ to 3’ direction. Sequence highlighted in **bold** represent restriction sites.

