## Supplemental Table 1 for "*Neisseria meningitidis* sibling small regulatory RNAs connect metabolism with colonization by controlling propionate use"

**Table S1. Strains and plasmids used in this study**

| **Strains or plasmids** | **Genotype or description** | **Reference** |
| --- | --- | --- |
| *N. meningitidis* |  | This study; unless indicated |
| H44/76 | Wild-type serogroup B strain | (1) |
| Δ*nmsRs* | H44/76 with *nmsRs* replaced by a Kan cassette, Kan^r^ |  |
| Δ*nmsRs* *gene*::*mcherry* + pEN11 | H44/76 Δ*nmsRs* *gene*::*mcherry* strain transformed with pEN11-*empty*, Kan^r^, Cm^r^ |  |
| Δ*nmsRs* *gene*::*mcherry* + pEN11_*nmsRs* | H44/76 Δ*nmsRs* *gene*::*mcherry* strain complemented with pEN11-*nmsRs,* Kan^r^, Cm^r^ |  |
| *prpB*::*mcherry* | Δ*nmsRs* derivative;  translational *prpB*::*mcherry* fusion, Cm^r^ |  |
| *prpC*::*mcherry* | Δ*nmsRs* derivative;  translational *prpC*::*mcherry* fusion, Cm^r^ |  |
| *prpF*::*mcherry* | Δ*nmsRs* derivative;  translational *prpF*::*mcherry* fusion, Cm^r^ |  |
| *ackA-*1::*mcherry* | Δ*nmsRs* derivative;  translational *ackA*1::*mcherry* fusion, Cm^r^ |  |
| *glyA*::*mcherry* | Δ*nmsRs* derivative;  translational *glyA*::*mcherry* fusion, Cm^r^ |  |
| *gatC*::*mcherry* | Δ*nmsRs* derivative;  translational *gatC*::*mcherry* fusion, Cm^r^ |  |
| *porA*::*mcherry* | Δ*nmsRs* derivative;  translational *porA*::*mcherry* fusion, Cm^r^ |  |
| *mmsB*::*mcherry* | Δ*nmsRs* derivative;  translational *mmsB*::*mcherry* fusion, Cm^r^ |  |
| *gdhR*::*mcherry* | Δ*nmsRs* derivative;  translational *gdhR*::*mcherry* fusion, Cm^r^ |  |
| *E.coli strains* |  |  |
| TOP10 | mcrA Δ(mrr-hsdRMS-mcrBC)  φ80lacZDM15 ΔlacX74 deoR  recA1 araD139 Δ(ara-leu)7697  galU galK rpsL endA1 nupG | Invitrogen |
| *Plasmids* |  |  |
| pMCG5 | pBluescript derivative;  allowing chromosomal expression of *mcherry* under control of the *pilE* promoter, Cm^r^ | Stratagene (2) |
| pMCG5_mut | pMCG5 derivative;  NsiI (+1) and NheI (ATG) restriction sites, Cm^r^ |  |
| pEN11-*pldA* | Cm^r^ and ErmC^r^ | (3) |
| pEN11 | pEN11-*pldA* derivative;  Δ*pldA,* Cm^r^ and ErmC^r^ | (4) |
| pEN11-*nmsRs* | pEN11-*pldA* derivative;  Δ*pldA*::*nmsRs,* Cm^r^ and ErmC^r^ |  |
