## Supplemental Table 2 for "*Neisseria meningitidis* sibling small regulatory RNAs connect metabolism with colonization by controlling propionate use"

**Table S2. Oligonucleotides used in this study**

| **Name** | **Sequence (5’→3’)^a^** | **Utilization** |
| --- | --- | --- |
| pMGC5**_**mut_NheI_F2 | GGAGTAATTTTATG**GCTAGC**AAGGGCGAGGAGG | Amplification of pMCG5 to introduce an NheI restriction site to generate pMCG5_mut |
| pMGC5_mut_NheI_R2 | CCTCCTCGCCCTT**GCTAGC**CATAAAATTACTCC |  |
| pMGC5**_**mut_NsiI_F2 | CCAATCTTGCTTTATAAT**ATGCAT**GTATCGCAACAAAATACC | Amplification of pMCG5 to introduce an NheI restriction site to generate pMCG5_mut |
| pMGC5_mut_NsiI_R2 | GGTATTTTGTTGCGATAC**ATGCAT**ATTATAAAGCAAGATTGG |  |
| SB_UTR_*prpC*_F | GTTTT**ATGCAT**CAACAAATCCGCATCGGTC | Amplification of the 5’ UTR and beginning of *prpC* (NMB0431) with NsiI/NheI ends for cloning in frame with *mcherry* in pMCG5_mut |
| SB_UTR_*prpC*_R | GTTTT**GCTAGC**GGATTTTTTAGGTTTGAGGGTCGG |  |
| SB_UTR_*prpF*_F | GTTTT**ATGCAT**TGGGCAGAGCCCACGCTACATAAGG | Amplification of the 5’ UTR and beginning of *prpF* (NMB0434) with NsiI/NheI ends for cloning in frame with *mcherry* in pMCG5_mut |
| SB_UTR_*prpF*_R | GTTTT**GCTAGC**TGTACCGCCACGGTAGTAAACGGC |  |
| SB_UTR_*ackA-*1_F | GTTTT**ATGCAT**GCCGTCTGAAACACCTTGCGC | Amplification of the 5’ UTR and beginning of *ackA*1 (NMB0435) with NsiI/NheI ends for cloning in frame with *mcherry* in pMCG5_mut |
| SB_UTR_*ackA-*1_R | GTTTT**GCTAGC**GCCTTTGAGCGATGAACTGCCGCA |  |
| SB_UTR_*glyA*_F | GTTTT**ATGCAT**CACAAAGGAGCCGAGAACATG | Amplification of the 5’ UTR and beginning of *glyA* (NMB1055) with NsiI/NheI ends for cloning in frame with *mcherry* in pMCG5_mut |
| SB_UTR_*glyA*_R | GTTTT**GCTAGC**TGCCAAATCGGGGTCGTATTG |  |
| SB_UTR_*gatC*_F | GTTTT**ATGCATA**ACGACGGTATCTTTACCATC | Amplification of the 5’ UTR and beginning of *gatC* (NMB1355) with NsiI/NheI ends for cloning in frame with *mcherry* in pMCG5_mut |
| SB_UTR_*gatC*_R | GTTTT**GCTAGC**GGAGAGTCGGGCGATTTTGTC |  |
| SB_UTR_*porA* _F | GTTTT**ATGCAT**GTATCGGGTGTTTGCCCGATG | Amplification of the 5’ UTR and beginning of *porA* (NMB1429) with NsiI/NheI ends for cloning in frame with *mcherry* in pMCG5_mut |
| SB_UTR_*porA* _R | GTTTT**GCTAGC**aagcggcagtgcggacaatac |  |
| SB_UTR_*mmsB*_F | GTTTT**ATGCAT**TACACTACAAACAGAAAAGG | Amplification of the 5’ UTR and beginning of *mmsB* (NMB1584) with NsiI/NheI ends for cloning in frame with *mcherry* in pMCG5_mut |
| SB_UTR_*mmsB*_R | GTTTT**GCTAGC**CATTTGCCCTAAGCCTATCCA |  |
| SB_UTR_*gdhR*_F | GTTTT**ATGCAT**GGAAGGGTTCTGCAGGGA | Amplification of the 5’ UTR and beginning of *gdhR* (NMB1711) with NsiI/NheI ends for cloning in frame with *mcherry* in pMCG5_mut |
| SB_UTR_*gdhR*_R | GTTTT**GCTAGC**CAATACCGACAATACCTGATC |  |
| SB_*nmsRs* _Kan_F2 | CAGCGACTGCAACCACAACTG | Amplification of the Kanamycine resistance cassette |
| SB_*nmsRs* _Kan_R2 | CAACAGATGGGGATTGAGATG |  |
| SB_con2_F | CCATGCCAATAGAGATACCCCACG | Forward primer on *recC* for verification of *gene*::*mcherry* constructs |
| SB_*mtrF*_R | CCCACATTCTATCCCGCACC | Reverse primer on *mtrF* for verification of *gene*::*mcherry* constructs |
| Copy_pEN11_F1 | CGACTACATCAACCGCAATGATAAATATTCAAAGCGGTTAGATGATTTAC | Amplification of pEN11_*nmsRs* |
| Copy_pEN11_R1 | GTAAATCATCTAACCGCTTTGAATATTTATCATTGCGGTTGATGTAGTCG |  |
| SB_qPCR _*prpB*_F | AGCCATGACCGATTTGAACA | Amplification of *prpB* by RT-qPCR |
| SB_qPCR _*prpB*_R | AAACTCGGTAATGTTCGCCA |  |
| SB_qPCR _*prpC*_F | CCAAGCTCAAATCCATGC | Amplification of *prpC* by RT-qPCR |
| SB_qPCR _*prpC*_R | CATCCGATGGACGTAATGCG |  |
| SB_qPCR_*prpF*_F | GAAGCGCACGCGACAAAATC | Amplification of *prpF* by RT-qPCR |
| SB_qPCR _*prpF*_R | GTCCGAACGCGCCGATCACG |  |
| SB_qPCR _*ackA*1_F | CACGACCGCATCAAAGCCATC | Amplification of *ackA-*1by RT-qPCR |
| SB_qPCR _*ackA*1_R | ACAGCGAGTCTGTTTTGATCG |  |
| SB_qPCR _*glyA*_F | GCGAATACGTCGATATTGTCG | Amplification of *glyA* by RT-qPCR |
| SB_qPCR _*glyA*_R | CGACACCATTTTGGGTATGTC |  |
| SB_qPCR _*gatC*_F | CCATTAACACAGACGGCATCG | Amplification of *gatC* by RT-qPCR |
| SB_qPCR _*gatC*_R | GAAGTACGCAACCGTCTGTAC |  |
| SB_qPCR _*porA* _F | ATGTGGCTTCGCAATTGG | Amplification of *porA* by RT-qPCR |
| SB_qPCR _*porA* _R | GTTTCAGCGGCAGCGTTC |  |
| SB_qPCR _*mmsB*_F | CTGATGGTTTCCGACTATGC | Amplification of *mmsB* by RT-qPCR |
| SB_qPCR _*mmsB*_R | CGTCAAAGCACTTGTCGAAG |  |
| SB_qPCR _*gdhR*_F | GCGAAAGCGGCAATTTGGAAC | Amplification of *gdhR* by RT-qPCR |
| SB_qPCR _*gdhR*_R | GCACAATTTGTTGTTCAGCC |  |
| SB_*rmpM*_F | ACAACCTGAAAGTATTGGCG | Amplification of *rmpM* by RT-qPCR |
| SB_*rmpM*_R | TCAGAACCCATAAAGTCGGT |  |
| SB_*ccba*_F | ACCGCATCAAAGAAATCCAC | Amplification of *cbbA* by RT-qPCR |
| SB_*ccba*_R | TGACTTTCAGCCATTCTTGC |  |
| SB_*nmsR_A_*_F2 | CCGTCGAGTTGCTTGA | Amplification of *nmsR_A_* by RT-qPCR |
| SB_ *nmsR_A_*_R3 | CCAGTTGAATGTGTGCCA |  |
| SB_ *nmsR_B_*_F | CGTTAGCTGGTTCGAGTAGTCAGTTAATAG | Amplification of *nmsR_B_* by RT-qPCR |
| SB_ *nmsR_B_*_R | CCAGTTGAATGTGTGCCAAGTCTACAAAGGAG |  |

a. Sequence highlighted in **bold** represent restriction sites
