## Supplementary figures and images for "*Neisseria meningitidis* sibling small regulatory RNAs connect metabolism with colonization by controlling propionate use"

### Supplemental Figure 1

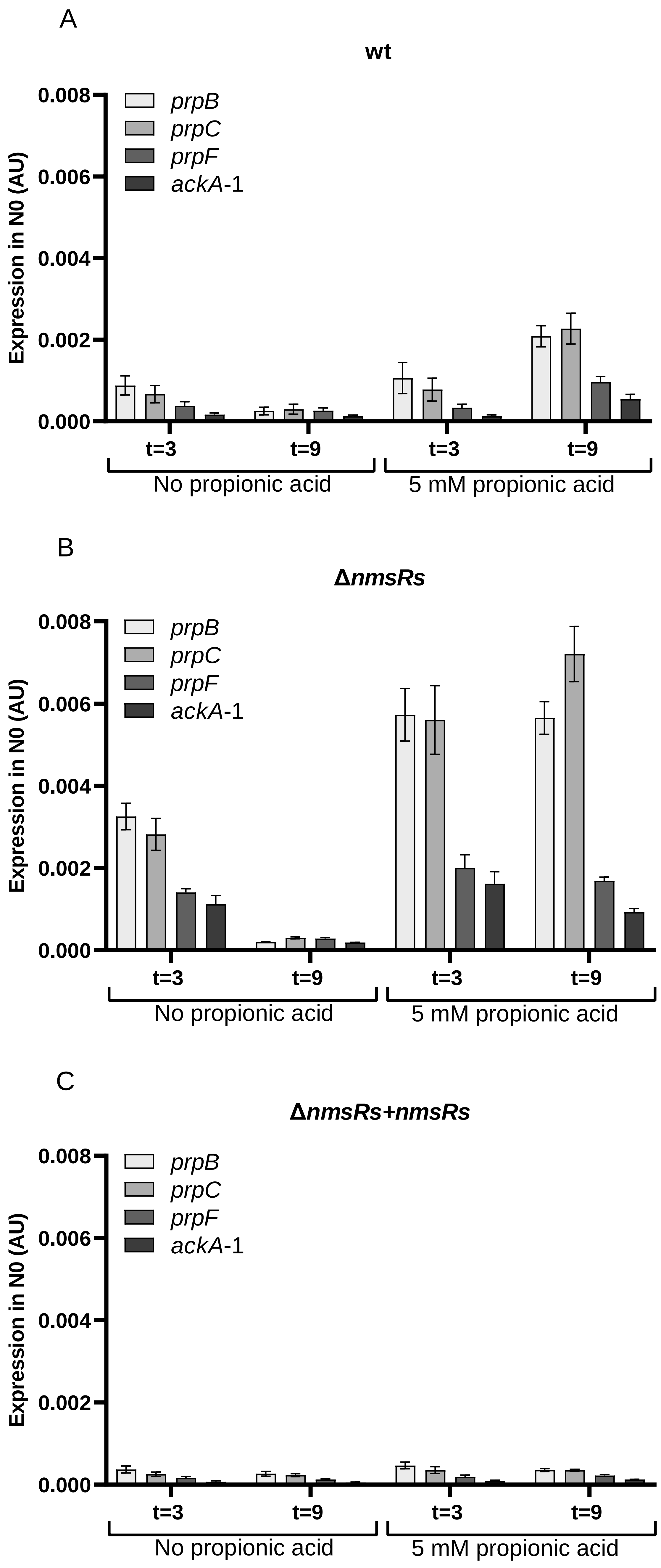

### Supplemental Figure 2

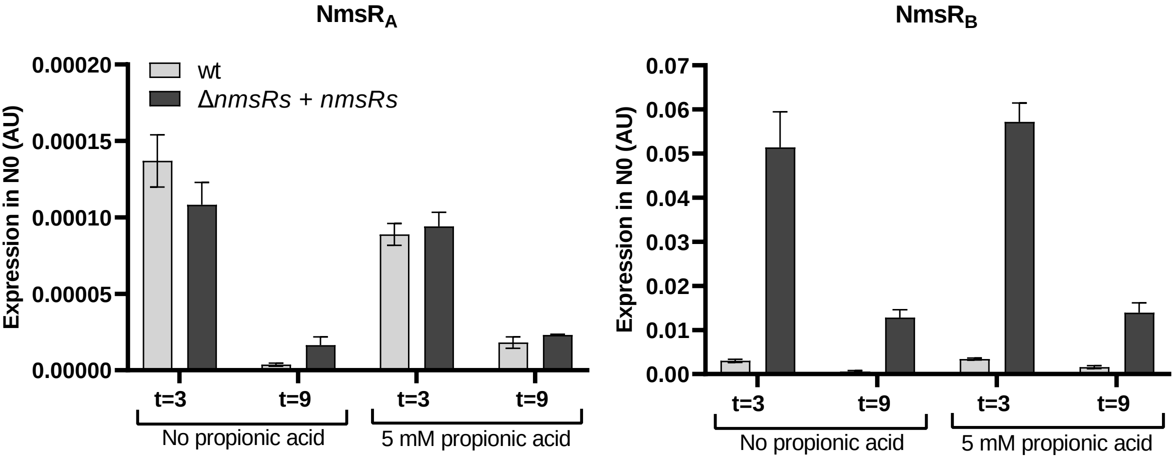
